# Dissociable roles of theta and alpha in sub-second and supra-second time reproduction: An investigation of their links to depression and anxiety

**DOI:** 10.1101/2022.02.14.480446

**Authors:** Mingli Liang, Sara Lomayesva, Eve A. Isham

## Abstract

The Striatal Beat Frequency model (Matell & Meck, 2004) suggests that multiple cortical oscillators contribute to human temporal cognition. Based on the theory, the current study examined the interactions between interval timing and neural oscillations, as predicted by timing deficits found in neuropsychiatric conditions. The interaction is predicted to manifest in frontal-midline theta and occipital alpha oscillations, which are two important cortical oscillatory sources engaged in working memory and time perception. Therefore, the main goal of the study is to examine how depression and anxiety modulate frontal-midline theta and occipital data during the encoding of sub- and supra-second intervals. In the current study, participants reproduced sub- (400, 600, 800ms) and supra-second (1600, 1800, 2000ms) intervals while they underwent scalp EEG recordings. Anxiety and depression levels were measured via self-report psychometrics. We found that higher levels of self-reported anxiety and depression were associated with shorter reproduction of lengths of durations. Further, time-frequency analysis of scalp EEG revealed a dissociation where state anxiety altered the links between frontal-midline theta power and sub-second interval reproduction, while depression and trait anxiety altered the association between occipital alpha power and supra-second interval reproduction. Our results suggest that anxiety and depression alter time perception, and this differentially relates to frontal-midline theta and occipital alpha as substrates for sub- and supra-second interval timing.

## 1. Introduction

### 1.1. Background

Neural oscillations, a prevalent signature of neuronal activity that can be recorded with scalp electroencephalogram (EEG), have been theorized and documented as potential neural substrates that support time perception. Two proposed neural mechanisms supporting interval timing is through coincidence detection (Striatal Beat Frequency model, Matell and Meck, 2004), and neural integration model (Simen et al., 2011), The coincidence detection model hypothesizes that multivariate patterns of cortical oscillators of various frequencies and brain regions contribute to interval timing. It is theorized that these cortical oscillators act as pacemakers and reset their phases at the onset of the to-be-encoded time interval, similar to a starting gun signal during a race (Kononowicz, 2015). Subsequently, patterns generated by these pacemakers are subjected to coincidence detection in the striatum by spiny neurons to determine differences in temporal intervals and estimate interval length. The neural integration model (for a review, see Paton & Buonomano, 2018; Simen et al., 2011)suggests that interval timing is supported by populations of neurons via leaky integration, are predicts that three types of cells are involved in interval timing: one that stays active during entire intervals, one that ramps up their activities (accumulator) during intervals, and one that activates at the interval timing thresholds. While both theories are built upon activities of multiunit spiking trains, it is not clear how neural oscillations during interval timing map to the theorical components in these two theories.

Two types of oscillations are robustly observed in scalp EEG and fit well as candidates for time cognition: theta oscillations (4-8Hz) in the frontal midline region, and alpha oscillations in the occipital region (8-12Hz). Frontal midline theta oscillations are typically observed with maximal amplitudes around electrode Fz, and at ~7 Hz (Mitchell et al., 2008). Midfrontal theta oscillations are recruited during interval timing, and the degree of midfrontal theta activation is diminished in the Parkinson Diseases group compared to healthy controls (Singh et al., 2021). Similar activation of theta oscillations in midfrontal cortices and striatum have also been observed in rodent models during interval timing behavior (Emmons et al., 2016; Gu et al., 2018). Increases of frontal midline theta oscillations have also been associated with natural locomotion (Liang et al., 2018), working memory performance (Hsieh & Ranganath, 2014; Karakaş, 2020), mental arithmetic calculation (Gärtner, Grimm, & Bajbouj, 2015), and in association with interval discrimination task (Liang et al., 2021). Past research has also suggested that alpha frequencies determine the time window for visual perception (Cecere, Rees, & Romei, 2015), and modulation of alpha frequencies causally change temporal discrimination performance (Mioni et al., 2020). Furthermore, the alpha frequency band has also been postulated as a cortical oscillator that could be capable of modulating the speed of the ‘internal clock’ (Treisman et al., 1994), therefore implying an additional role of alpha as an indicator of temporal expectation (Praamstra et al., 2006; Cravo et al., 2011). Such a marker may further serve as a selective coordinator for temporal prioritization of sensory events (Jensen et al, 2014; Kononowicz & van Wassenhove, 2016). Modulation of alpha frequencies via transcranial alternating current stimulation has also been shown to directly cause change in temporal discrimination performance (Mioni et al., 2020).

Studying individuals diagnosed with neuropsychiatric disorders associated with clear differences in behavioral working memory capacity (e.g., attention deficit hyperactivity disorder, see Ptacek et al., 2019 for review) or neural activation in the striatum (e.g., Parkinson’s disorder, Harrington et al., 2011) has been a crucial approach toward validating the SBF model (Allman & Meck, 2012). Individuals experiencing depression or anxiety, two of the most prominent behavioral health conditions in the United States (U.S. Department of Health and Human Services, 2022), often perform poorly on tasks involving working memory and attention (Christopher & MacDonald, 2005; Moran, 2016), though mixed results have been found with respect to interval timing tasks. Some studies have shown that those living with depression have the tendency to produce shorter time durations compared to controls (Gil & Droit-Volet, 2009). Other studies have found no effect of depression symptomatology on time perception (e.g., Oberfeld, Thones, Palayoor, & Hecht, 2014). Individuals with anxiety have been shown to provide longer reproduced intervals than the actual target interval, especially in fearful situations (Bar-Haim et al., 2010). Recent work suggests this might be modulated by attention (Liu & Li, 2020); that is, attentional bias toward negative stimuli could alter the influence of state anxiety on interval timing. Further still, a study comparing time reproduction in clinical anxiety and depression patients found that both types of patients had the tendency to under-reproduce three different temporal intervals, though anxious patients were more likely to under-reproduce to a greater degree than depressed patients (Mioni et al., 2016).

Beyond the correlation between mental health and time perception, various associations among self-reported mental health symptoms and neural oscillations have also been reported. It has been observed that posterior alpha power is negatively related to depression levels (Jiang, et al., 2016 in adults; Umemoto et al., 2021 reported in adolescents). Furthermore, it has been proposed that alpha oscillations in the posterior regions contribute to the modulation of top-down control or inhibition (Jensen and Mazaheri, 2010). Such perspective has been linked to attention and memory (Haegens, Händel, & Jensen, 2011; Jiang et al., 2015b). In addition to alpha oscillations, theta oscillations have also been used as a biomarker for anxiety. In animal models, mice lacking serotonin display higher levels of anxious behaviors also exhibit a larger increase in theta power in the hippocampus (Gordon et al., 2005; see Okonogi & Sasaki, 2021 for review). Given the evidence of associations, albeit inconclusive, between neural activity associated with timing and mental health characteristics, a closer look into how these neural oscillations behave during timing tasks in anxious and depressed individuals is needed.

### 1.2 Current study

The central aim of the current study is to observe activities of specific neural oscillations during interval timing and to evaluate whether these neural activities differ between individuals experiencing distinct mental health-related issues. To further explore the contributions of cortical oscillators in the encoding and reproduction of temporal information, we examined the correlational relationship between depression and anxiety level and the strength of oscillatory activities during encoding of sub- and supra-second intervals. We focused on frontal-midline theta and occipital alpha oscillations based on the established associations with temporal cognition and mental health. The goal is to provide further insight into how cortical oscillations may serve as a biomarker for the relationship between interval timing and mental health, as predicted by the SBF model (Matell & Meck, 2004).

## 2. Materials and methods

### 2.1. Participants

Fifty undergraduate students at the University of Arizona (31 females, 19 males, age range between 18 to 32 years old, with a mean age of 19.9) participated in this study. They received either monetary compensation or course credit. All participants provided informed consent and the experiment protocol was approved by University of Arizona Institutional Review Board.

### 2.2. Materials and Procedure

Participants were seated approximately 60 cm away from a computer monitor and were tested individually. They were asked to complete a brief demographic questionnaire in addition to three psychometric measures of anxiety and depression symptomatology (described below) before beginning the time reproduction task. SuperLab 6 software (Cedrus Corporation, San Pedro, CA) was used to create the experimental stimuli presentation. The refresh rate of the monitor is 60Hz. In each experimental trial, a black stick figure appeared at the center of the screen on a plain white background for a brief duration of either 400, 600, 800, 1600, 1800, or 2000 ms (the order was randomized across trials). Immediately upon the offset of the stimulus, the participants reproduced the interval by pressing down on the spacebar twice: once to indicate the beginning and once more to mark the end of the remembered duration. In doing so, the same black stick figure appeared in the center of the screen as a visual placeholder. Participants were instructed to try their best to remember the duration they were presented with for each trial and were encouraged to refrain from counting or using physical means of timekeeping (e.g., tapping) during the encoding phase. Each participant completed 10 trials per duration, yielding a total of 60 trials. Data from the 400, 600 and 800 ms trials were grouped as sub-second timescale and those from the 1600, 1800 and 2000 ms were classified as supra-second timescale. Each experimental session ranged from 60 to 90 minutes.

### 2.3. Self-report anxiety and depression measures

Participants completed the Beck Depression Inventory-II (BDI-II; Beck, Steer, & Brown, 1996) and the State-Trait Anxiety Inventory (STAI; Spielberger et al., 1983) to assess degrees of abnormality in current mental health state. The BDI-II measures the presence of depression symptoms based on established clinical diagnostic materials (American Psychiatric Association, 1994) via a 21-item self-report rating inventory. The higher the BDI scores, the greater the degree of depression. The STAI has two forms that measure two main types of anxiety: state (STAI-S) and trait anxiety (STAI-T). State anxiety refers to the level of anxiety being experienced in the present moment, while trait anxiety refers to a personality predisposition to perceive situations as threatening (Speilberger et al., 1983).

### 2.4. EEG data acquisition and preprocessing

Continuous EEG data was recorded at a sampling rate of 2500Hz with a 64-channel BrainVision actiCHamp system (BrainVision LLC, Morrisville, NC). Electrodes were placed on the scalp according to the international 10-20 system. The reference electrode was FCz. Impedances of electrodes were reduced to below 5kΩ prior to starting the recording. Event markers for the onset and offset of the time interval encoding period, as well as the participant spacebar presses indicating the time reproduction start and end, were recorded on the continuous EEG data via a Cedrus c-pod device (Cedrus Corporation, San Pedro, CA).

Preprocessing and analyses were performed with EEGLAB (Delorme & Makeig, 2004) in MATLAB and MNE in Python. An average referencing scheme was used offline, and a 2-50Hz bandpass filter was applied to the continuous data. Independent component analysis (ICA) was performed to ameliorate eye-blinks, eye-movements, and muscle movement artifacts. After ICA artifact rejection, data was downsampled to 250Hz.

### 2.5. Power spectra

To compute the power spectra, we used 6-cycle Morlet wavelets, sampling 15 frequency steps logarithmically from 2 to 30Hz. Continuous data was epoched surrounding the onsets of standard duration encoding, including data from two seconds before the onsets and four seconds after the onsets. The epoched data was used to compute power spectra. Time-frequency power from one second after the encoding onsets was averaged to represent the encoding activities for sub-second trials (400ms, 600ms and 800ms). For supra-second trials (1.6s, 1.8s, and 2s), timefrequency power was averaged from 1.6 seconds, 1.8 seconds, and 2 seconds after the encoding onsets, respectively, to represent the encoding activities.

### 2.6. Onset-related Event-related Potentials (ERPs) and Evoked Time-frequency Analyses

To calculate the ERPs time-locked to the onsets of encoding, we averaged the raw traces from electrodes Fz and POz across all trials within a condition, and averaged them across 51 participants. To examine the spectral contents of the ERPs, we performed a timefrequency analysis of the subject-level averaged ERPs. Time-frequency power of the ERPs were extracted via 6-cycle Morlet Wavelets using MNE package. Oscillatory power between 2Hz to 64Hz were extracted in 30 logarithmic steps (the following frequencies were covered: 2, 2.25, 2.54, 2.86, 3.23, 3.64, 4.1, 4.62, 5.2, 5.86, 6.61, 7.45, 8.39, 9.46, 10.66, 12.01, 13.53, 15.25, 17.19, 19.37, 21.83, 24.6, 27.72, 31.24, 35.21, 39.68, 44.72, 50.39, 56.79, and 64 Hz. The oscillatory power were log10 transformed and zscored within each subject for each frequency. The time-frequency power matrices from 51 subjects were then submitted to a cluster-based permutation test to evaluate their statistical significance (Maris & Oostenveld, 2007) using 500 permutations and a t-critcal value of 1.656.

### 2.7. Code and data accessibility

Deidentified scalp EEG data and analyses codes are available upon request. Part of the analysis code is openly available on GitHub: https://github.com/liangmingli/mental_health_time_cognition.

## 3. Results

### 3.1. Participants reproduced time as a function of the standard durations

We first analyzed whether participants successfully learned the standard durations and reproduced them as instructed. Results suggested that indeed participants showed above chance performance, reproducing time as a function of the standard durations (Fig. 1A). When averaged across all responses, participants reproduced 400ms standard durations as 616.88±509.24ms, 600ms durations as 812.40±916.63ms, 800ms durations as 965.88±558.70ms, 1600ms durations as 1630.74±520.18ms, 1800ms durations as 1829.91±830.81ms, and 2000ms durations as 2014.07±932.09ms. We also normalized the time reproduction responses as a ratio to the standard durations, and again participants showed above chance performances reproducing the intervals (Fig. 1B): for 400ms, they reproduced with an averaged ratio of 1.54±1.27; for 600ms, 1.35±1.53; for 800ms, 1.21±0.70; for 1600ms, 1.02±0.33; for 1800ms, 1.02±0.46; and for 2000ms durations, 1.01±0.47. The pattern of these ratios follows Vierordt’s law (Fortin & Rousseau, 1998) such that shorter intervals have a greater tendency to be overestimated.

**Figure 1.**
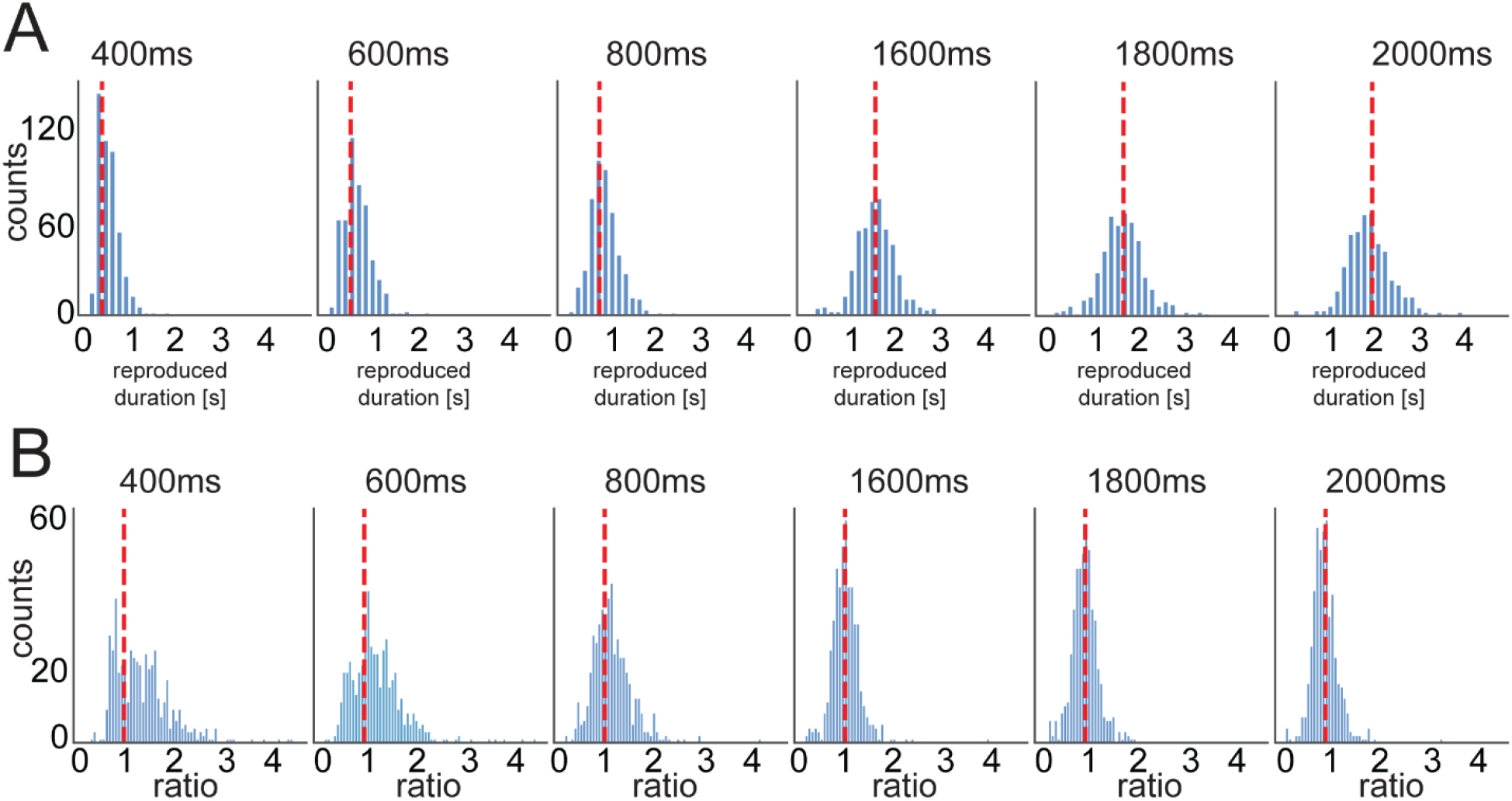
Time reproduction performance as a function of the standard durations. (**A**) Histogram of trial-wise time reproduction responses. When averaged, participants accurately reproduced both sub-second and supra-second intervals. Red lines indicate the positions of standard durations. (**B**) Histogram of normalized time reproduction responses. Normalization was achieved by ratios (dividing raw reproduction responses by standard durations). A ratio of 1 means a perfect reproduction of standard durations. For visualization purposes, responses longer than 5 times the standard durations were not displayed. Red lines indicate the positions of perfect time reproduction (i.e., ratio = 1).

### 3.2. Frontal midline theta and occipital alpha oscillations were recruited during both sub-second and supra-second time encoding

Frontal midline theta and occipital alpha are two prominent oscillatory signatures in human scalp EEG, and they have been demonstrated to be relevant to sensory processing, learning and memory, and decision-making. Here, we report an observation of frontal-midline theta and occipital alpha oscillations during the encoding periods for both sub-second and suprasecond durations (Figs. 2 A, B). We focus specifically on the power spectra of electrode sites Fz and POz (Figs. 2 C, D): Fz displays maximum amplitudes of frontal midline theta oscillations (Mitchell et al., 2008), and POz is at a location where salient occipital alpha oscillations are typically observed (Benwell et al., 2019; Sutterer et al., 2021), respectively. Visual examination of both spectra revealed an increase in power around 6 Hz (theta) in the electrode Fz, and around 10 Hz (alpha) at the electrode POz. When analyzing the event-related potentials (ERPs) at frontal electrode Fz, clear oscillations in theta range were visible (Figs. 1 E and F for 400ms condition and 2000ms condition). Time-frequency analyses of the ERPs across 51 participants revealed an evoked activation of theta (4-8Hz) oscillations at electrode Fz across all intervals (see Figs.1 E and F, and supplementary figure Fig S1). These findings suggest that during the encoding periods of standard durations, both frontal-midline theta and occipital alpha oscillations were recruited. In the next set of analyses, we examined how frontal midline theta and occipital alpha activity during encoding were related to the time reproduction outputs, and how such relationships were modulated by anxiety and depression levels among our participants.

**Figure 2.**
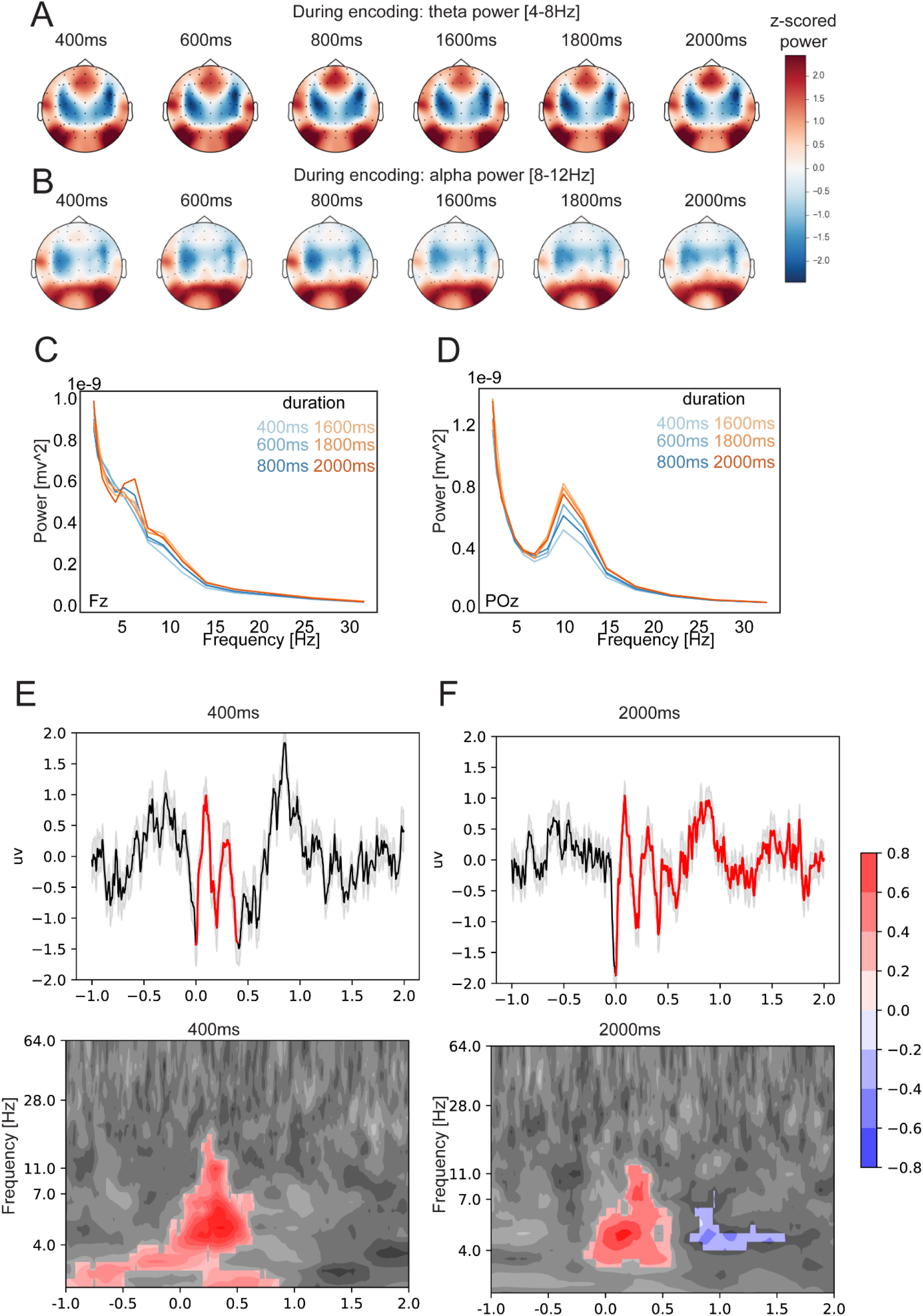
Frontal midline theta and occipital alpha oscillations were recruited during both subsecond and supra-second duration encoding. (**A**) Topography plots of z-scored theta power averaged across the standard duration encoding windows and averaged across participants. During both sub- and supra-second duration encoding, theta power peaked near electrode Fz with a frontal-midline topography. (**B**) Topography plots of z-scored alpha power averaged across the encoding windows and participants. During both sub- and supra-second encoding, alpha power was concentrated near the occipital region. (**C**) Power spectra at electrode Fz averaged across participants. An increase in power around the theta frequency range (4-8Hz) is visible in the averaged power spectra during duration encoding. (**D**) Power spectra at electrode POz averaged across participants. An increase in power around the alpha frequency range (8-12Hz) is salient during encoding for all standard durations (400ms −2000ms). (**E) (F**) Encoding-onset-related ERPs are visible in both sub- (**E, 400ms**) and supra-second (**F, 2000ms**) conditions at midfrontal electrode Fz. ERPs were obtained by averaging across trials and 51 participants. Time-frequency analyses performed on subject-specific ERP revealed a phase-locked evoked component of theta (4-8Hz) oscillations at midfrontal electrode Fz. The top rows of subplots present the onset-related ERPs, shades indicate standard errors across 51 participants, and the red lines indicate the standard-interval encoding periods. The bottom rows of subplots present the time-frequency analyses of the onset-related ERPs, and the values presented are z-scored power. Red and blue clusters indicate statistically significant (*p* < .05) clusters returned by permutation clustering algorithms (Maris & Oostenveld, 2007). For a complete list of ERPs and related time-frequency plots across all durations, please see supplementary Figure S1.

### 3.3. Greater degree of depression and anxiety correlated with shorter reproduction

Given the mixed evidence of the effects of anxiety and depression symptomology on timing performance, we analyzed how the lengths of interval reproductions were tied to the selfreport depression level (BDI-II), state anxiety level (STAI-S), and trait anxiety level (STAI-T). We median-split the self-report depression and anxiety scores into low/high anxiety and low/high depression groups, and we subjected the reproduced intervals to three separate 2 (low vs high symptom presence; between subjects) x 2 (sub-second vs supra-second durations) mixed ANOVAs according to depression, state, and trait anxiety (Fig. 3). The ANOVA results demonstrated that higher levels of self-reported depression and anxiety were associated with shorter reproduced temporal intervals (main effect for depression, *F*(1, 296) = 3.30, *p* = .007; main effect for state anxiety, *F*(1, 296) = 5.27, *p* = .002; main effect for trait anxiety, *F*(1,296) = 16.77, *p* = 5.44e-5). In addition, by treating the psychometric scores as continuous variables, we evaluated multiple linear regression models, using the standard durations (600ms-2000ms) and the psychometric scores as predictor variables to predict time reproduction performance. The results from regression models corroborated the similar conclusion that the higher the level of self-reported depression, the shorter the reproduced interval (linear regression, *t*(297) = −2.60, *p* = .01, R^2^ = 0.27); and the higher the level of self-reported state anxiety, the shorter the interval reproduction (linear regression, *t*(297) = −3.68, *p* < .001, R^2^ = 0.29); and lastly, the higher the level of self-reported trait anxiety, the shorter the interval reproduction responses (linear regression, *t*(297) = −3.34, *p* = .001, R^2^ = 0.28). Together, these findings suggest that self-report levels of anxiety and depression were negatively related to the length and fidelity of interval reproduction responses.

**Figure 3.**
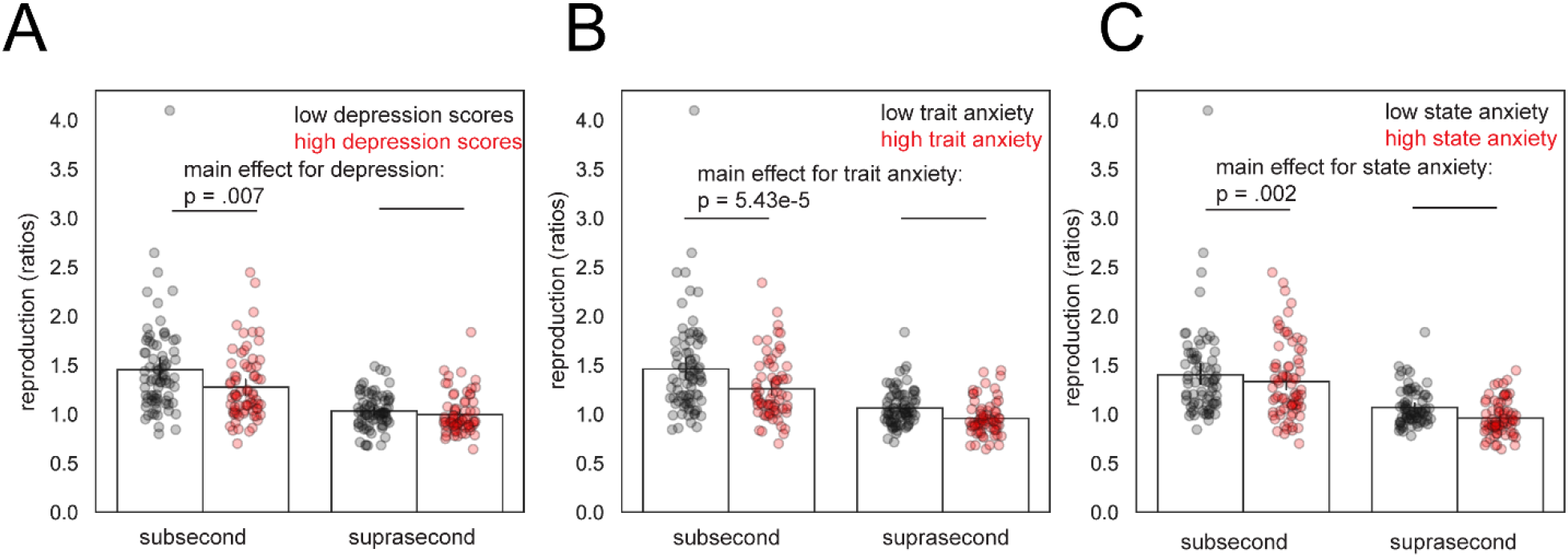
Anxiety and depression levels negatively correlated with reproduction response. Time reproduction ratios were subjected to a 2 timescales (sub-second and supra-second) x 2 mental health median-split (low and high; between-subjects) mixed ANOVA. (**A**) Higher depression scores are linked to shorter time reproduction (*p* = .007). (**B**) Higher trait anxiety scores are associated with shorter time reproduction (*p* = 5.43e-5). (**C**) Higher state anxiety scores are correlated with shorter time reproduction (*p* = .002). Notes: error bars indicate 95% errors. The reproduction ratios in the sub-second timescale were higher than the ratio in the supra-second timescale, consistent with Vierordt’s law (Fortin & Rousseau, 1998).

### 3.4. Anxiety modulated the correlations between theta power and the reproduction of sub-second intervals

We next examined whether frontal-midline theta and occipital alpha oscillations varied with depression and anxiety levels, and whether such changes were associated with reproduction performance. Previous research suggests that temporal cognition for sub- and supra-second time scales involve distinct neural circuits (Mauk & Bunomono, 2004) and are engaged in different sensory and cognitive functions (see Szelag, Stanczyk, & Szymaszek, 2022 for review). Therefore, we dichotomized our data as sub- and supra-second intervals and analyzed them accordingly.

To address whether depression and anxiety affect the association between theta and alpha and the reproduction of sub-second intervals, we calculated the correlations between theta and alpha power and the lengths of reproduction. The returned statistics, in the form of t statistics indexed the correlational strengths. Subsequently, we examined how these correlational strengths were linked to the level of self-report depression and anxiety. The data revealed that anxiety scores were positively associated with the theta correlational strengths (Fig. 4A, *t* = 2.07, *p* = .04, linear regression). No associations were found between the correlational strengths and depression scores (*t* = 0.05, *p* = .96), nor between the correlational strengths and trait anxiety scores (*t* = 0.04, *p* = .97). Additionally, no significant links were observed between the alpha correlational strengths and depression (*t* = −0.73, *p* = .47), state anxiety scores (*t* = −1.05, *p* = .30), nor trait anxiety scores (*t* = −1.31, *p* = .20). Collectively, these findings imply that the level of state anxiety modulates the links between frontal midline theta and sub-second time reproduction.

**Figure 4.**
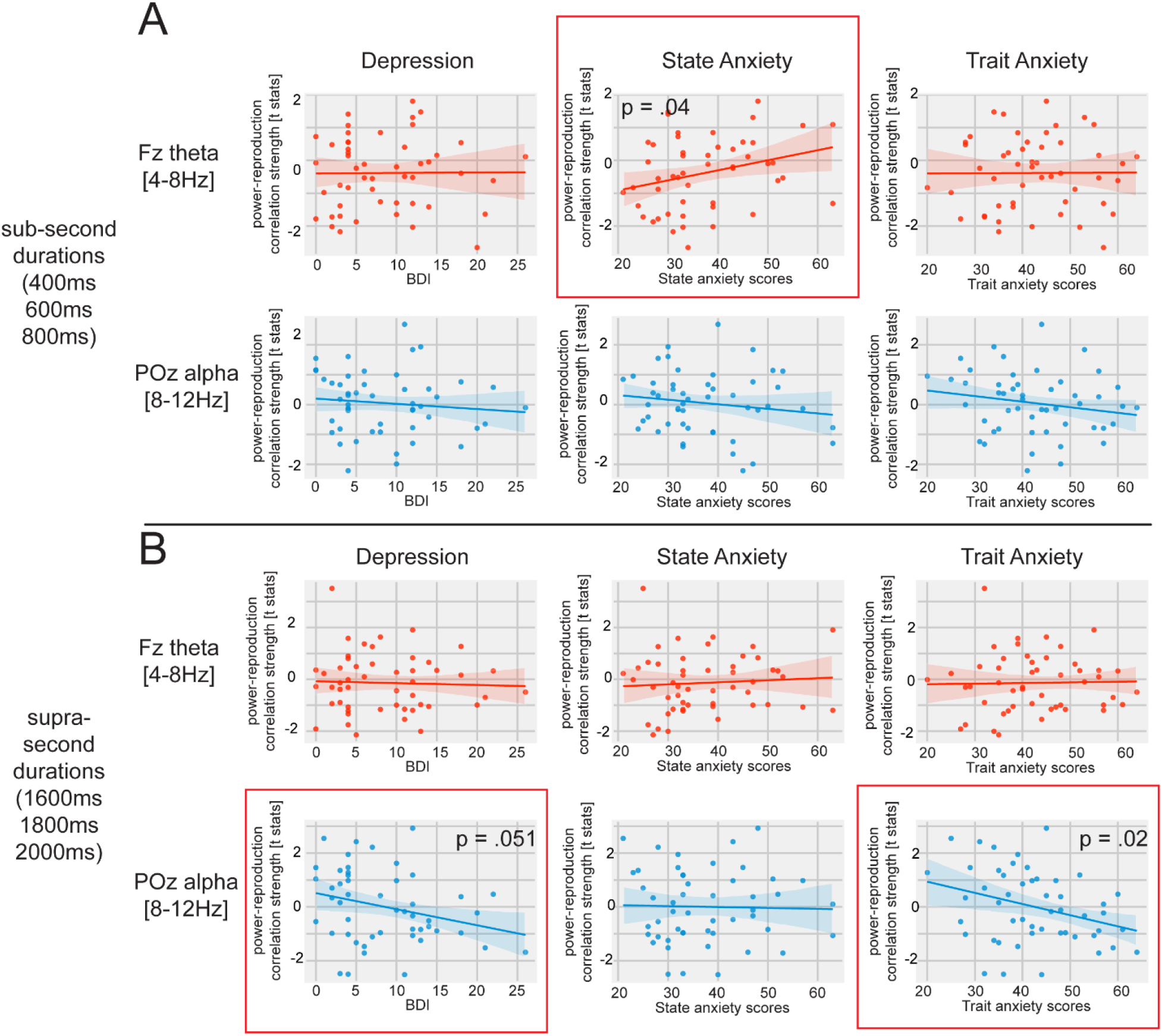
Depression and anxiety modulated the relationships between time reproduction and neural oscillations. (**A**) State anxiety scores significantly modulated the association between Fz theta power and sub-second time reproduction (*p* = .04). Such modulation in the sub-second durations were not found with other psychometric variables or with occipital alpha power (*p* >.05). (**B**) Trait anxiety and depression modulated the association between occipital alpha power and supra-second time reproduction (*p* = .02, *p* = .05, respectively). Such modulation was not found in Fz theta power or in state anxiety (*p* > .05). Note: Red boxes indicate significance or trending significance of modulation.

### 3.5. Depression and anxiety modulated the correlations between alpha power and the reproduction of supra-second intervals

We next examined whether depression and anxiety altered the links between theta and alpha oscillations and the reproduction of supra-second intervals. Our data demonstrated that selfreport trait anxiety level was negatively associated with the alpha power correlational strengths

(Fig. 4B, *t* = −2.32, *p* = .02). Self-reported depression symptoms showed a similar negative association with a trending significance (Fig. 4B, *t* = −2.00, *p* = .051), and self-reported state anxiety level did not show modulation of the alpha power correlational strengths (*t* = −0.18, *p* = .86). No modulation on the theta power correlational strengths was observed with respect to depression (*t* = −0.28, *p* =.78), state anxiety (*t* = 0.50, *p* = .62), nor the trait anxiety (*t* = 0.16, *p* = .87). These observations, therefore, suggest that trait anxiety and depression may serve a moderating role on the link between occipital alpha and the reproduction of supra-second intervals.

## 4. Discussion

The current study examined how neural oscillatory activities, namely frontal theta and occipital alpha oscillations, underlie the changes of temporal interval reproduction performance accompanied by varying degrees of self-reported anxiety and depression symptoms. Firstly, at the behavioral level, we observed that individuals who scored higher on the BDI-II, indexing a higher degree of depression, generally reproduced shorter durations than those who scored on the lower end. This observation of an under reproduction is consistent with past reports for individuals with depression (Gil and Droit-Volet, 2009). We also found that those with higher trait or state anxiety, just like with depression, also under reproduced the target intervals, which is consistent with previous observations of performance based on clinical diagnosis (Mioni et al., 2016), but inconsistent with performance proceeded by fear induction (e.g., Bar-Haim, et al., 2010).

The high comorbidity of anxiety and depression symptomatology could potentially explain why participants with either high trait anxiety or depression had the tendency to reproduce intervals that were shorter than the standard interval (Gorman, 1996; Mineka, Watson, & Clark, 1998). This comorbidity has been thought to be rooted in a feedback loop of rumination; for example, depression associated with past events contributes to anxiety and worry of what the future may hold, impacting the ability to function in the present. Some of the participants in the current study had both high anxiety and high depression; it is possible that either more dominant etiology (e.g., depressed mood or anxious arousal), or the combined effect of co-occurring depression and anxiety symptoms, contributed to the observed under reproduction performance. Unfortunately, this speculation could not be verified with the current study; because of the quasi-experimental design we were not able to classify the participants systematically or sufficiently to meet the statistical power. Nevertheless, we believe our findings will help pave ways for a future study that examines how time perception may be affected by comorbid clinical neuropsychological traits.

In addition to behavioral performance, we also observed the involvement of frontal-midline theta and occipital alpha oscillations across different timescales and across different levels of neuropsychological conditions. This is consistent with existing findings showing that midfrontal oscillations emerge as a responsive signal to conflicts, punishments and errors, and more anxious individuals show greater activation of midfrontal theta during those scenarios (Cavanagh & Shackman, 2015). Specifically, in the sub-second timescale, but not in the suprasecond range, state anxiety scores positively correlated with the association between frontal midline theta power and sub-second reproduction. Although correlation does not equal causation, we speculate that the significant correlation could be interpreted as that the direction between frontal-midline theta power and reproduction length flipped: those with higher state anxiety levels demonstrated a positive association between theta power and sub-second reproduction lengths, while those with lower state anxiety level demonstrated a negative association between theta power and sub-second reproduction. On top of the findings by Cavanagh and Shackman (2015) that midfrontal theta is activated after feedbacks, our findings suggest that midfrontal theta oscillations are also recruited in the context of interval timing, before feedback is given, and that such activated is modulated by self-reported anxiety/depression. This indicates that midfrontal theta might regulate multiple facets of interval timing, spanning from encoding, retrieval, and post-retrieval evaluation. We speculate that the observed relationship between frontal theta power, anxiety level, and the sub-second interval reproduction reflects more neural resources being dedicated to the encoding process of temporal information as state anxiety levels increase. This can be partially explained by the fact that more attentional resources are needed to process brief intervals in the sub-second range. This interpretation is consistent with previous reports demonstrating that encoding of durations involve slow frequency oscillations such as delta-theta oscillations (Gu & Meck, 2015; Matell & Meck, 2004), and consistent with the role of frontal-midline theta oscillations in memory encoding and mental calculation (Gärtner et al., 2015; Hsieh & Ranganath, 2014). We emphasize here that our findings with respect to theta and alpha activity might not represent the processing of durations alone, but instead might also include how much attention and effort are needed. As such, this interpretation aligns with the theoretical perspective that oscillatory patterns may capture both interval timing and working memory (Gu & Meck, 2015).

Occipital alpha oscillations likely serve a different role in the encoding of supra-second durations, distinct from the role of frontal-midline theta oscillations. In our investigation, we observed a dissociation between frontal-midline theta and occipital alpha oscillations when accounting for reproduction performance and mental health symptoms. Specifically, for those with higher scores of trait anxiety or depression, the correlation between occipital alpha power and reproduction lengths tended to be negative. Conversely, for those with lower scores of trait anxiety or depression, the correlation between alpha power and reproduction tended to be positive. Our results support the pacemaker theory that alpha band modulates the speed of the internal clock via working memory capacity (Treisman et al., 1994). As such, the results also suggest that alpha frequencies are the cortical oscillators proposed in the SBF model (Matell & Meck, 2004). Higher alpha power may imply a faster internal clock speed, which in turn, would lead to a greater accumulation of temporal information and therefore the perception of a longer duration. The presence of trait anxiety or depression symptoms may fundamentally alter how the internal clock contributes to the temporal cognition network.

Our findings support the notion that sub-second and supra-second interval reproductions involve different neural substrates (Mauk & Buonomano, 2004), and that anxiety and depression symptoms differentially impact the neural substrates for temporal cognition. While we assume both sub- and supra-second interval timing involve the encoding and memory of the target durations, each timescale could alter the neural activity in the fronto-central cortical projections in different ways. Sub-second intervals, for instance, are associated with automatic processes such as timing processes (e.g., reflexes) that are too short for an individual to fully consciously experience but will be recognized after the interval has passed via memory. Supra-second durations, on the other hand, are involved in more effortful processes (e.g., memory) and are more accessible to be recognized in the present conscious moment (White, 2017). As such, the approach that an organism takes toward encoding, along with the expectation on how to use the temporal information, may be different, and this may give rise to the differences in the neural oscillatory responses associated with sub- and supra-second timing observed in the current study.

One limitation of our study is that depression and anxiety scores show collinearity in our dataset. To study the differential effects of anxiety and depression on interval timing behavior, it would be ideal to examine a pool of participants with dissociated levels of anxiety and depression. Given the limitations, we interpreted our data separately for anxiety and depression with caution, according to each analysis. Future investigations could attempt to collect a larger sample size to cover population with non-comorbidity in depression and anxiety.

While our findings support important roles of frontal-midline theta and occipital alpha oscillations along with how they are related to effects of depression and anxiety on interval timing, other neural oscillations of distinct frequency bands should also be considered. For instance, beta and gamma frequency bands are involved in working memory and are potential markers for depression (Fitzgerald & Watson, 2018). Beta rhythms, possibly a variant of Rolandic mu rhythm (Caplan, et al., 2003) in particular, are interpreted to be involved in working memory (Tallon-Baudry, 2003) especially for time estimation (Wiener et al., 2018), along with sensorimotor maintenance (Engle & Fries, 2010) and motor learning in anxiety (Sporn et al., 2020). Future research may also consider if resting state oscillatory asymmetry found in depressed and anxious individuals could contribute to differences in interval timing compared to controls (see Thibodeau, Jorgensen & Kim, 2006 for review).

## Acknowledgments

This research was supported by the University of Arizona startup funds for EI. We thank Matthew Grilli for serving as our mental health consultant, and Arne Ekstrom for valuable feedback on an earlier version of the manuscript.

