## Supplementary material for "Dissociable roles of theta and alpha in sub-second and supra-second time reproduction: An investigation of their links to depression and anxiety": supp.fig.1

Mingli Liang^1^

Sara Lomayesva^1^

Eve A. Isham^1,2^

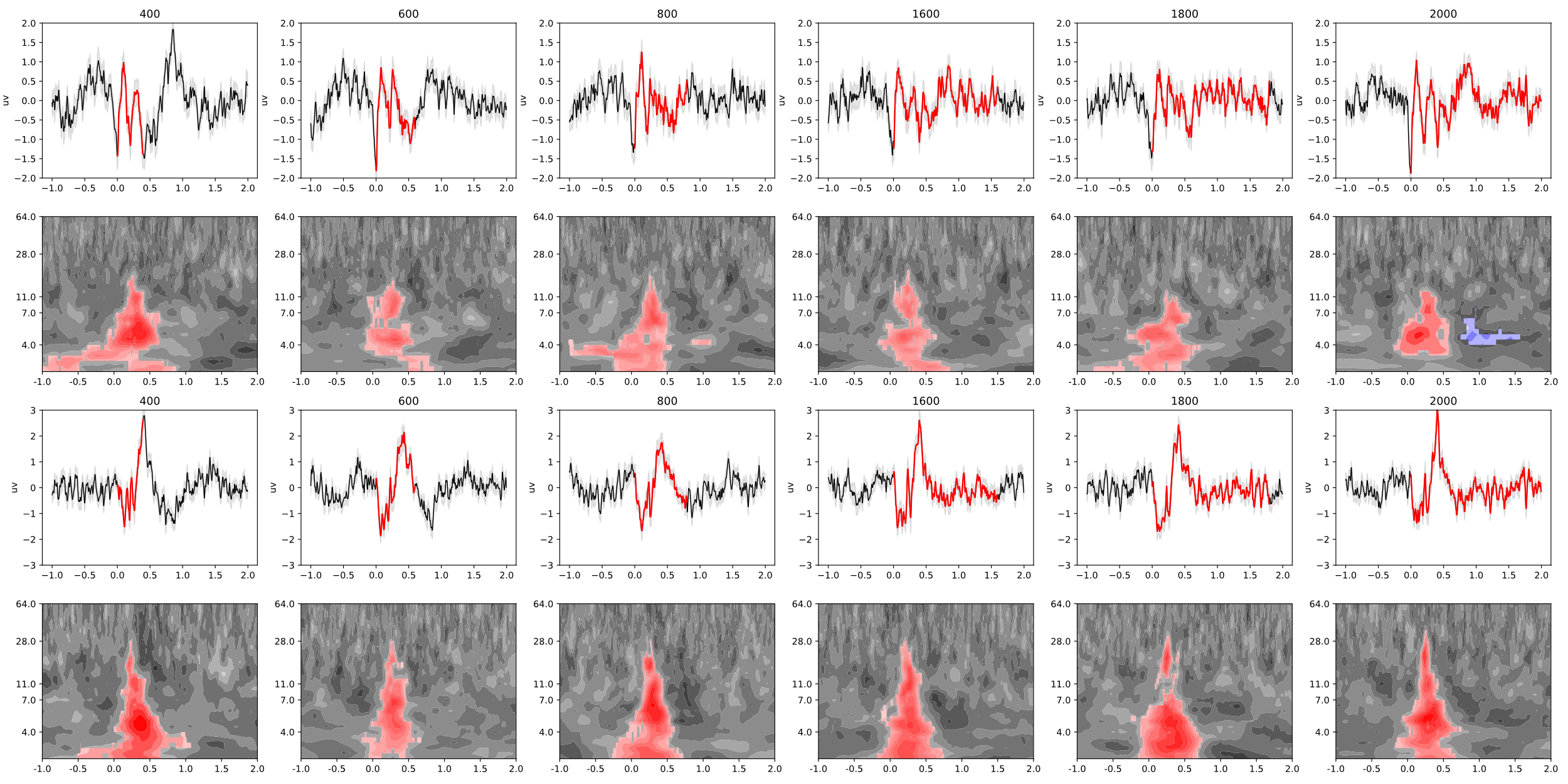

**Figure S1.** Encoding-onset-related ERPs are visible in both sub- and supra-second conditions at midfrontal electrode Fz and posterior electrode POz. The **top two** rows show data at electrode **Fz**, the ERPs and evoked time-frequency analyses. The **bottom two** rows show data from electrode **POz**. ERPs were obtained by averaging across trials and 51 participants. Time-frequency analyses performed on subject-specific ERP revealed a phase-locked evoked component of theta (4-8Hz) oscillations at midfrontal electrode Fz, across all six possible intervals (indicated by the significant clusters returned by permutation tests). At electrode POz, large amplitude ERPs are observed with the peaks near 500ms post encoding onsets. Time-frequency analyses of the ERPs at POz revealed a significant broadband power increase covering 2-30Hz.

*Notes*: Shades in the ERP subplots indicate standard errors across 51 participants, and the red lines indicate the standard-interval encoding periods. The color values in the time-frequency plots presents z-scored power. Red and blue clusters indicate statistically significant (*p* < .05) clusters returned by permutation clustering algorithms (Maris & Oostenveld, 2007).
